# Learning Conjunctive Representations

**DOI:** 10.1101/2024.05.30.596595

**Authors:** Markus Pettersen, Frederik Rogge, Mikkel Elle Lepperød

## Abstract

Hippocampal place cells are known for their spatially selective firing patterns, which has led to the suggestion that they encode an animal’s location. However, place cells also respond to contextual cues, such as smell. Furthermore, they have the ability to remap, wherein the firing fields and rates of cells change in response to environmental changes. How place cell responses emerge, and how these representations remap is not fully understood. In this work, we propose a similarity-based objective function that translates proximity in space, to proximity in representation. We show that a neural network trained to minimize the proposed objective learns place-like representations. We also show that the proposed objective is trivially extended to include other sources of information, such as context information, in the same way. When trained to encode multiple contexts, networks learn distinct representations, exhibiting remapping behaviors between contexts. The proposed objective is invariant to orthogonal transformations. Such transformations of the original trained representation (e.g. rotations), therefore yield new representations distinct from the original, without explicit relearning, akin to remapping. Our findings shed new light on the formation and encoding properties of place cells, and also demonstrate an interesting case of representational reuse.

## 1 Introduction

Animals and humans are capable of extraordinary feats of navigation, from birds migrating by following magnetic fields [Packmor et al., 2021], to rodents navigating mazes [Small, 1901] and cab drivers memorizing and traversing nearly 26000 busy London streets [Fernandez-Velasco and Spiers, 2023]. In the brain, navigation ability is believed to be supported by Hippocampal place cells [O’Keefe and Dostrovsky, 1971, O’Keefe and Nadel, 1978]. Place cells are known for their tendency to only fire at one, or a few locations within a recording environment [Park et al., 2011], correlating with the position of the animal. Besides encoding allocentric location, place cells also respond to other contextual cues, such as room identity, room geometry, odors or colors [Latuske et al., 2018, O’Keefe and Burgess, 1996, Leutgeb et al., 2005b, Jeffery, 2011]. In other words, place cells form conjunctive representations, merging spatial information with available contextual cues. It is also believed that place cells distinguish contextual information trough so-called remapping. In response to large changes in the environment, place cell responses change, with cell responses becoming uncorrelated across contexts [Muller and Kubie, 1987, Leutgeb et al., 2004, 2005b].

How place cells obtain their striking behaviors remains a matter of debate. Some argue that place cells inherit their firing fields from upstream cell types, such as grid cells [Hafting et al., 2005, Solstad et al., 2006, Jeffery, 2011], or border cells [Barry et al., 2006, Pettersen et al., 2024] or that place cell representations in different environments form distinct attractor states [Jeffery, 2011, Samsonovich and McNaughton, 1997]. However, exactly how place-like representations emerge, and how they can be learned, remains poorly understood.

Recently, a range of normative models have demonstrated that neural networks trained to solve simple navigation tasks actually learn representations similar to biological spatial cells [Cueva and Wei, 2018, Banino et al., 2018, Sorscher et al., 2022, Whittington et al., 2020, Xu et al., 2022, Dorrell et al., 2022, Schaeffer et al., 2023]. However, these models often feature complicated architectures and a range of different regularization strategies, making it difficult to discern why the observed representations actually arise. Furthermore, these works tend to focus on learning grid cell-like representations, often placing less emphasis on other emergent cell types, such as place cells.

In this work, we take inspiration from existing machine learning models [Cueva and Wei, 2018, Banino et al., 2018, Sorscher et al., 2022, Xu et al., 2022, Dorrell et al., 2022] and propose a minimal, similarity-based objective. When a feedforward network is trained to minimize this objective, we find that it learns place-like spatial representations. We further show that the objective is easily extended to encompass conjunctive representations of space and context. When trained in a conjunctive setting across multiple contexts, we find that the network learns uncorrelated representations for different contexts, similar to Hippocampal global remapping. We further train a recurrent neural network to solve the same task, and find that band cell-like [Krupic et al., 2012] representations emerge alongside the place code, showing that other cell types may be involved in path integration, and also that the objective extends to more naturalistic settings. Lastly, we show that we can apply orthogonal transformations to learned spatial representations in order to generate new spatial maps while preserving the similarity structure. Thus, the proposed objective allows for switching between different maps without having to relearn them, offering an interesting perspective on Hippocampal remapping.

## 2 Methods

### 2.1 Proposed objective function

First, we tackle the problem of learning an encoding of some region of physical space (e.g. a square recording enclosure). We follow the example set by recent normative models [Schaeffer et al., 2023, Dorrell et al., 2022], and argue that biologically plausible spatial representations can be obtained directly by specifying the properties of the network population vector. Considering properties that a spatial representation should obey, we propose an objective where we demand that: 1) Points that are close in physical space, should be represented by similar population vectors. 2) Points that are distant in physical space, should be represented by dissimilar population vectors. 3) In the open field, no point or direction is special, so the above properties should be rotation- and translation invariant. 4) Unit activations are bounded. 5) Unit activations are non-negative.

To investigate representations with these properties, we suggest training neural networks to minimize a spatial encoding objective. Consider a neural network with population vector **p**(**z**_*t*_), where each component *p*_*i*_(**z**_*t*_), *i* = 1, 2, 3, …, *N* is the firing rate of a particular (output) unit of the network at a particular location, **x**_*t*_ (where t indexes e.g. time along a trajectory or sampled spatial locations). **z**_*t*_, on the other hand, is the network’s latent position estimate corresponding to **x**_*t*_. We impose non-negativity in the architecture of the neural network, by selecting appropriate activation functions. In this work, we consider the ReLU. With these prerequisites in mind, we explore the following objective function

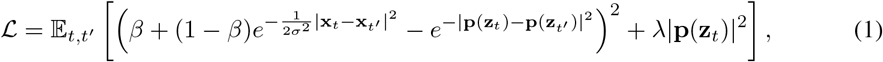

where |*·*| denotes the L2-norm, while *β* is a lower bound on the target similarity, *σ* is a hyperparameter that controls the scale of the learned similarity structure and *λ* a hyperparameter governing an L2 activity regularization term.

Intuitively, (1) compares the (Gaussian) similarity of two points x_*t*_ and x_*t*′_, and asserts that the corresponding population vectors at those points (**p**(**z**_*t*_) and **p**(**z**_*t*′_)) should exhibit the same similarity. In other words: Points that are close in physical space should be represented similarly, while distant points in physical space should be represented using dissimilar population vectors. This idea is illustrated in Fig. 1a). Notably, the Gaussian similarity is also translation invariant, and rotationally symmetric. Thus, every point and every direction adheres to the same “encoding rule”. The scaling term given by *β* ensures that dissimilar population vectors can still have overlapping unit activity, while still being highly different. This approach is similar in spirit to the concept of vectors being *nearly* orthogonal in hyperdimensional computing [Kanerva, 2009], which can greatly increases the capacity of a single vector to store information. When neural networks are trained to minimize (1), we find that individual units display spatial selectivity similar to biological place cells.

**Figure 1.**
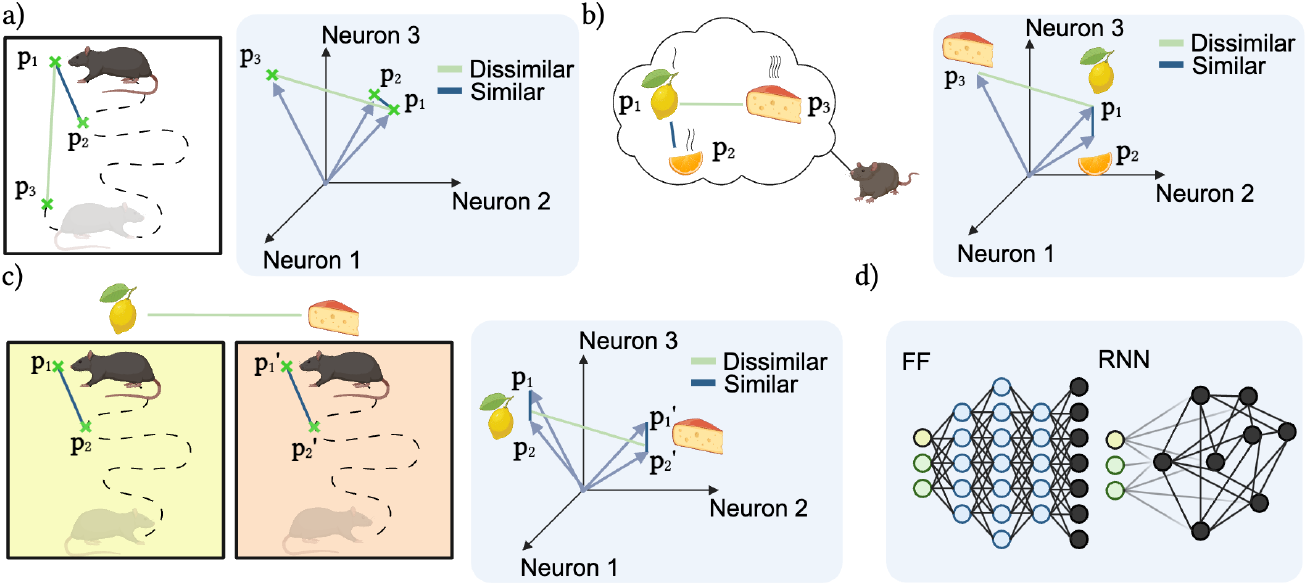
Overview of models and objective. a) Illustration of the spatial objective: locations that are close should be encoded by similar population vectors, distant locations by dissimilar population vectors. b) Similar to a); similar context signals should be represented by similar population vectors, dissimilar contexts by dissimilar population vectors. c) Similar to a) and b), but for conjunctive encoding of space and context. d) The explored neural network architectures; a feedforward, and a recurrent network.

We also find that the objective in (1) is just a special case of a more general encoding objective, where distinct sources of nonspatial information can be encoded in a single population vector. Fig. 1b) illustrates how contextual signals can be represented in a similar manner to the spatial case. However, place cells do not encode contextual information exclusively, but rather respond to particular locations and contexts in conjunction. Again, the similarity objective can be extended to accomodate this observation. If we represent context information in the simplest manner, as a scalar signal *c*, we can have the neural network encode spatial and contextual information conjunctively by training it to minimize

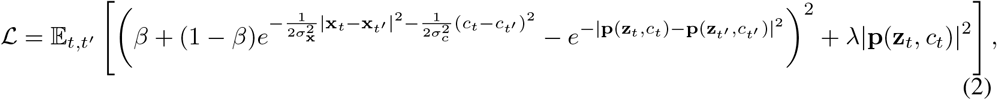

where, in general *σ*_x_ and *σ*_*c*_ are spatial and contextual encoding scales, respectively. For this work, however, we set *σ*_x_ = *σ*_*c*_. As with the purely spatial and purely contextual case, we model distinct spatial/contextual combinations as similar or dissimilar population vectors. An illustration of this situation is provided in Fig. 1c). We will see that networks trained to minimize (2) display a range of behaviors associated with place cells, including remapping.

### 2.2 Models and Training Details

In this work, we trained two distinct networks to minimize the proposed objective functions (1) and (2): a feedforward network and a recurrent network, as illustrated in Fig. 1d). Models were implemented and trained using the Pytorch library [Paszke et al., 2019].

The feedforward network featured two densely connected layers with 64 and 128 hidden units, followed by an output layer containing *n*_*p*_ = 100 units. Every layer was equipped with the ReLU activation function, ensuring non-negativity. Notably, the output units of the network together form the representation that is used to compute the loss, i.e. **p**. The weights of the feedforward network were all initialized according to input size-dependent uniform distribution, following the PyTorch default [Paszke et al., 2019].

For the spatial objective (1), the input to the feedforward network consisted of minibatches of Cartesian coordinates, sampled randomly and uniformly within a 2 *×* 2 square enclosure. For the conjunctive objective (2), the input to the network was a concatenation of randomly sampled Cartesian coordinates **x**, and uniformly sampled scalar context signals *c*, i.e. **input** = cat(**x**, *c*). Context signals were sampled uniformly in the interval *c* ∈ [−2, 2]. To increase the number of training samples, distances in either objective function were computed between minibatch elements.

The recurrent network consisted of a single vanilla recurrent layer equipped with the ReLU activation function, without added bias. Similar to the feedforward network, this network featured *n*_*p*_ = 100 recurrent units. In the spatial case, the recurrent state at a particular time *t* was given by

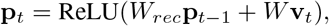

where 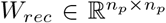 is a recurrent weight matrix, 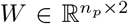 an input weight matrix and **v**_*t*_ Cartesian velocity inputs. In the case of conjunctive encoding, the input at time *t* was a concatenation of the velocity and a (time-constant) scalar context signal. Note that this also adds an additional column to the input weight matrix. To mitigate vanishing and exploding gradients, the recurrent weight matrix was initialized to the identity, similar to Le et al. [2015]. Similar to the feedforward network, losses were computed by comparing across trajectories (and contexts). The trajectory length was taken to be *T* = 10 timesteps. The initial recurrent state was computed by feeding the trajectory starting location (and optionally, the context) into a three-layer, densely connected network with 64, 64 and *n*_*p*_ = 100 units, respectively, each equipped with a ReLU activation function.

All networks were trained for a total of 50000 training steps, with a batch size of 64. Note that each training minibatch featured only previously unseen data, due to the ease with which new data could be generated (see section 2.4 for details). For each model, we used the Adam optimizer [Kingma and Ba, 2017] with a learning rate of 10^−4^.

### 2.3 Spatial Correlation & Remapping

To evaluate representational changes in the face of changing context input, we ran the trained feedforward network on 32 linearly spaced contexts (in the range [-2, 2]). Following Leutgeb et al. [2004], we then computed the spatial correlation between unit ratemaps across contexts. Between-context spatial correlations were computed by correlating the ratemap of a unit in one context with the same unit’s ratemap in another context in a binwise fashion. Note that ratemaps were only compared if a unit displayed non-zero activity in both contexts.

### 2.4 Dataset

The input to the feedforward network consisted of Cartesian coordinates, sampled in a 2 *×* 2 square enclosure. For the recurrent network, inputs consisted of velocity vectors along trajectories in the same square region. Trajectories were generated by creating boundary-avoiding trajectory steps successively. A step was formed by sampling heading directions according to a von Mises distribution, alongside step sizes drawn from a Rayleigh distribution. If the step landed the trajectory outside the enclosure, the velocity component normal to the wall was reversed, bouncing the trajectory off the boundary. Initial trajectory steps were sampled uniformly and randomly within the square arena. The von Mises scale was taken to be 4*π*, and the Rayleigh scale parameter 0.025. At training time, data was created on the fly due to the low computational cost.

### 2.5 Orthogonal Transformations

To explore how the spatial map of one context, 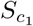, can be transformed into the spatial map of another context, 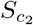, without explicit learning, we applied orthogonal transformations. These transformations preserve pairwise distances and norms, and the learned similarity structure across contexts. Our goal was to identify an orthogonal transformation *Q* that minimizes the Frobenius norm ∥ *·* ∥_*F*_ between the transformed spatial map from context *c*_1_ and the actual spatial map in context *c*_2_. Here, 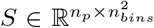 and the number of spatial bins *n*_*bins*_ was chosen to be 128. This problem is also known as the Procrustes orthogonal problem [Schönemann, 1966], which is formally defined as

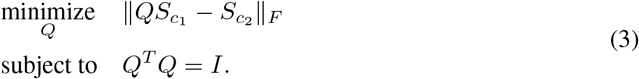

*Q* can be found by computing the Singular Value Decompositon of 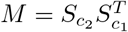 given by

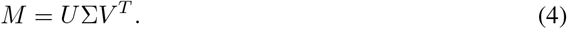

The orthogonal matrix *Q* can then be computed using *Q* = *UV* ^*T*^.

To explore whether orthogonal transformations of learned representation could be used to generate *new* representations (similar to remapping), we used gradient descent to *learn* an *n*_*p*_ *× n*_*p*_ nearorthogonal transformation matrix *Q*. While orthogonal matrices can be generated quite easily, we wanted to preserve the non-negativity of the place-like representation. We therefore minimized the following loss function for ratemaps of the feedforward network activity:

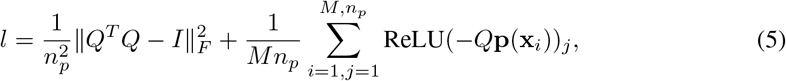

where *M* = 4096 and **x**_*i*_ indexes a regular grid of spatial points. Note that the final term amounts to a soft non-negativity constraint on the transformed representation. *Q* was trained for 10000 iterations, using Pytorch and the Adam optimizer with a learning rate of 10^−2^, and initialized according to a uniform distribution *𝒰* (−0.01, 0.01).

### 2.6 Decoding

In our model, each output unit is presumed to encode a spatial location near its peak activity. To determine the decoded position at a specific location *x*, we employed a weighted average of peak positions of the output units. The weights correspond to the activity levels of the respective units at that location. The decoded position 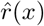 is thus given by:

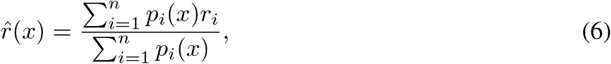

where 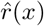 denotes the decoded position at *x, p*_*i*_(*x*) represents the activity of unit *i* at position *x*, and *r*_*i*_ indicates the peak location encoded by unit *i*. Notably, the decoding process does not incorporate all output units. Instead, units are prioritized based on their activity at position *x*, and only the top *n* most active units are included in the decoding.

### 2.7 Low-dimensional Projection

In order to investigate whether the population encoded lower-dimensional structures across different contexts, we employed Uniform Manifold Approximation and Projection (UMAP) [McInnes et al., 2018], with default parameters. We reduced the dimensionality of the data down to a two-dimensional space, and applied UMAP in two ways: first, to a collection of trajectories aggregated across various contexts, and second, to spatial maps within individual context.

### 2.8 Figures

Figures were created using BioRender.com.

### 2.9 Code Availability

Code to reproduce models and findings is available at https://github.com/bioAI-Oslo/ConjunctiveRepresentations.

## 3 Results & Discussion

### 3.1 Feedforward networks learn place cell-like representations

Training a feedforward network to minimize the spatial objective function (2), results in the emergence of place-like representations in the output units. These representations are analogous to place cells observed in the Hippocampus [O’Keefe and Dostrovsky, 1971], as illustrated by the example ratemaps in Fig. 2a). The parameter *σ*, which determines the width of the Gaussian kernel in the objective, directly influences the width of these learned representations. Notably, with sufficiently small values of *σ*, some units exhibit multiple place fields, akin to findings reported in Park et al. [2011]. The effectiveness of our model in learning the desired similarity structure imposed by the objective is evidenced by the saturated loss depicted in Fig. 2b) and the correspondence between the similarity of different points and the objective, as shown in Fig. 2f).

**Figure 2.**
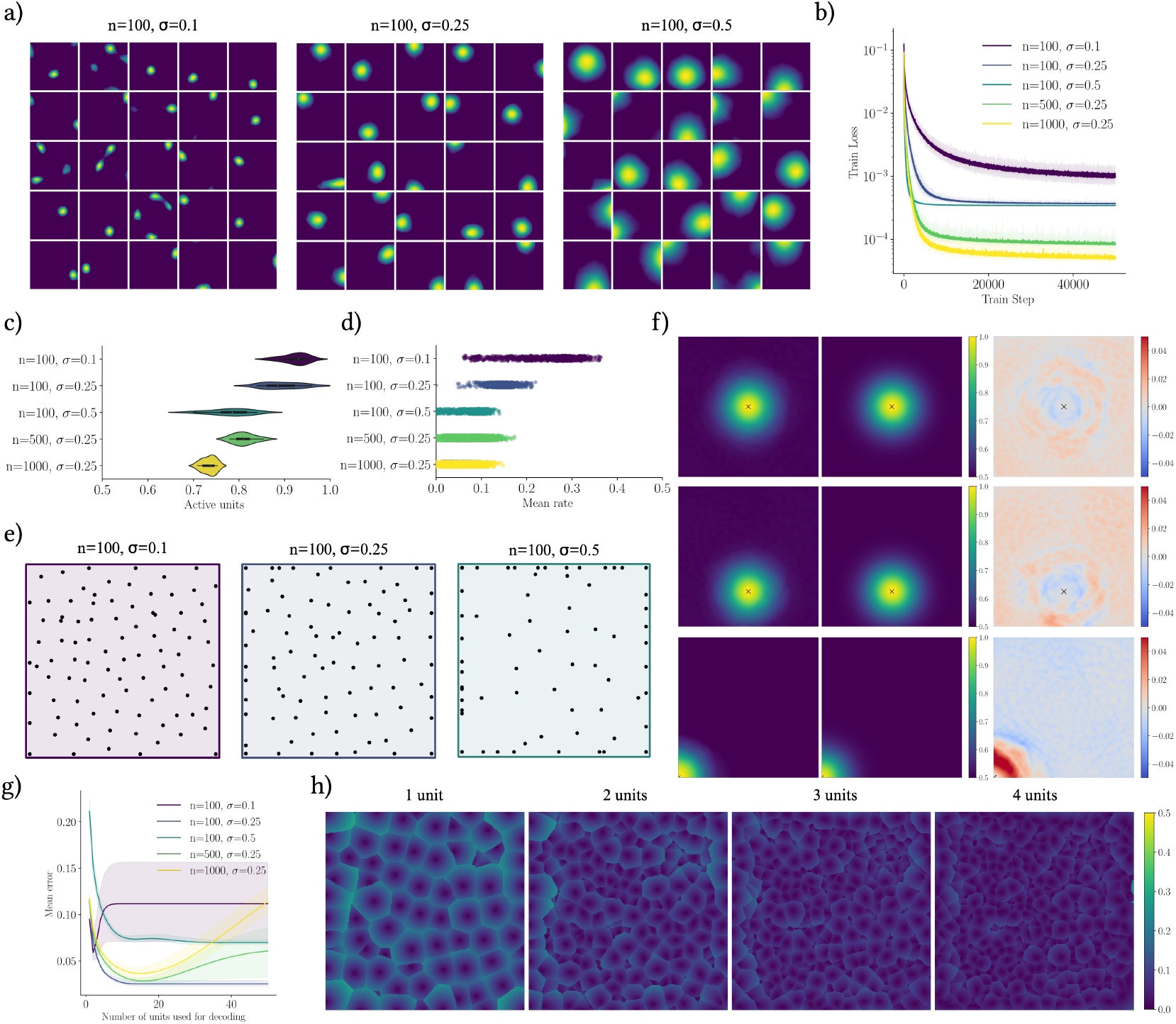
Feedforward network results. Ratemap examples of randomly selected active units for models with different values of *σ. n* is the number of output units. b) Training loss for different parameter combinations. Line shows the mean of 10 models. Error band shows the min and max across those 10 models. c) Distributions of the proportion of active units (mean rate *>* 0) for different parameter combinations. d) Distribution of mean rate of units pooled from all 10 models in each parameter combination. e) Spatial distribution of peaks of all units for different parameter combinations. f) Similarity structure for different points for the learned representations (left) and the objective (center), as well as the difference between the two (right). g) Mean position decoding error (measured by the euclidean distance) as a function of the number of units used for decoding. h) Decoding error maps for a varying number of units used for decoding; shown are results for models with *n*_*p*_ = 100 units and *σ* = 0.25.

Similar to biological place cells, the peak locations of our units span the entire training arena, as shown in Fig. 2e). Additionally, units display varied activity levels, as seen in Fig. 2d). Interestingly, not all units exhibit place-like tuning curves. Depending on the number of units *n*_*p*_ and the Gaussian kernel width *σ*, a subset of the units remains inactive across the entire arena, mirroring the behavior of place cell ensembles where some cells may not be active in certain conditions [Alme et al., 2014].

Next, we investigated whether the learned representations in our model indeed constitute effective spatial representations. To this end, we decoded the position based on (6), varying the number of units included in the analysis. The results, presented in Fig. 2g) and h), indicate that the decoding performance (measured by the mean Euclidean distance between the actual and decoded positions) differs based on the number of units utilized and the model configuration, and show that the learned representations can be used for accurate position decoding (achieving a mean error of a few percentage points relative to the arena size). These findings hint that there are optimal configurations of place field placement, size and unit count for decoding, providing valuable insights into the scaling of biological place cells.

### 3.2 Feedforward networks learn global-type remapping

When trained to encode multiple contexts in conjunction with spatial location, feedforward units exhibit dramatic firing field shifts, when comparing across contexts. This effect is shown in Fig. 3a), where ratemaps of randomly selected units are tracked while varying the context signal.

**Figure 3.**
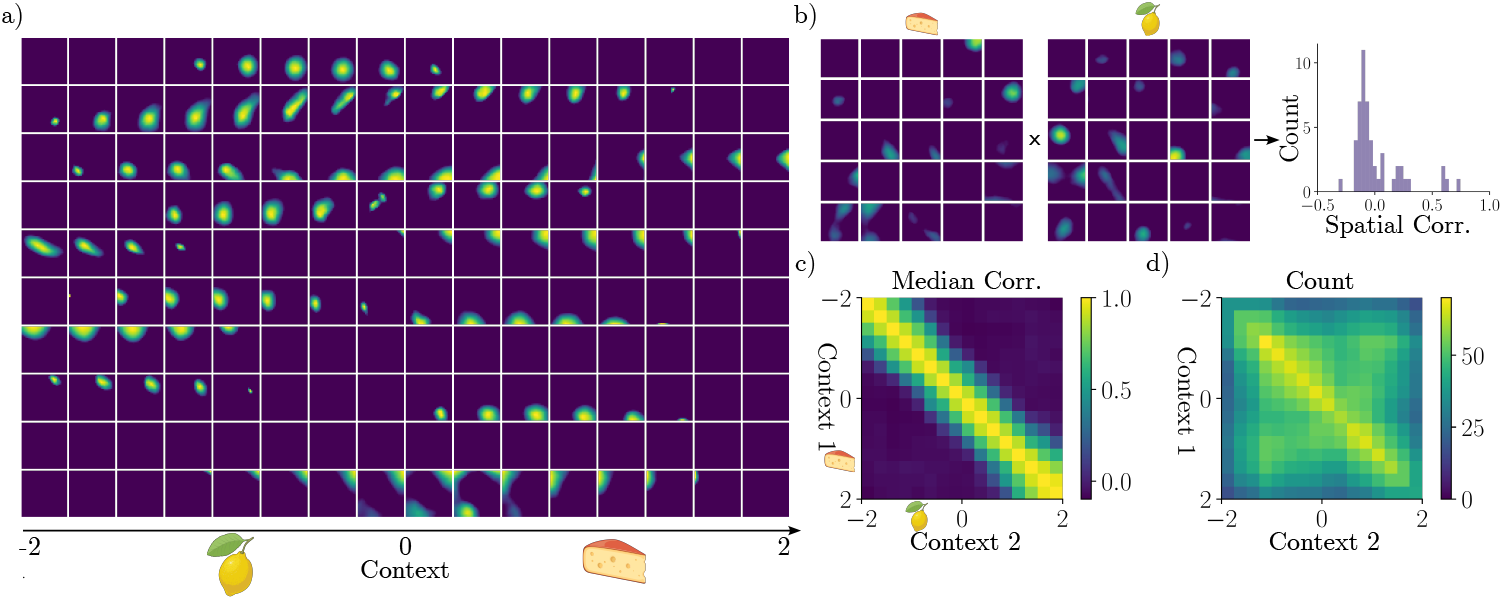
Feedforward network remapping results. a) Ratemaps as a function of context, for a random selection of 10 units. Each row tracks one unit. b) Example distribution of spatial correlations for ratemaps corresponding to two distinct contexts (context 1 =-0.9, context 2 = 1.2). c) Median spatial correlation when comparing across all contexts. d) Number of units included (units active in both contexts) in the analysis in c).

Notably, firing fields exhibit several place-like behaviors, such as multiple firing fields [Park et al., 2011] (e.g. bottom row), and field shifts between contexts, i.e. remapping [Muller and Kubie, 1987]. However, the observed remapping is not a sudden, attractor-like shift, which has been observed in place cells [Wills et al., 2005], but rather a gradual transition between states, which has also been seen during Hippocampal remapping [Leutgeb et al., 2005a].

Besides *appearing* similar to Hippocampal remapping, we find that spatial representations are indeed uncorrelated across different contexts, an example of which is shown in Fig. 3b). In fact, for sufficiently different contexts, the median spatial correlation between active units tends to zero, which aligns with global remapping in biological place cells [Leutgeb et al., 2004] (see Fig. 3c) and d)). Spatial correlations in Fig. 3c) also support the observation that units remap gradually, as correlations decay away from the diagonal.

### 3.3 Recurrent networks learn place- and band-like representations

When we train a recurrent network to solve the proposed objective function (1) while path integrating along simulated trajectories, we find that its units display place-like and band-like spatial tuning. Example ratemaps and trajectories are shown in Fig. 4a) and b), respectively. We also find that the recurrent network performs on par with the feedforward network, achieving similar loss minima, both for spatial and conjunctive encoding (see Fig. 4c), suggesting that the network has learned to path integrate. As band-like representations emerge only in the recurrent network, we speculate that these representations may be involved with path integration. Also worth noting is that while mean rates are similar, peak rates are markedly different between unit types (Fig. 4d) and e)), further hinting at different functional roles. That band cells contribute to path integration, has also been found in other neural network models [Schøyen et al., 2023]. As with the feedforward model, recurrent responses accurately capture the desired similarity structure (Fig. 4f)).

**Figure 4.**
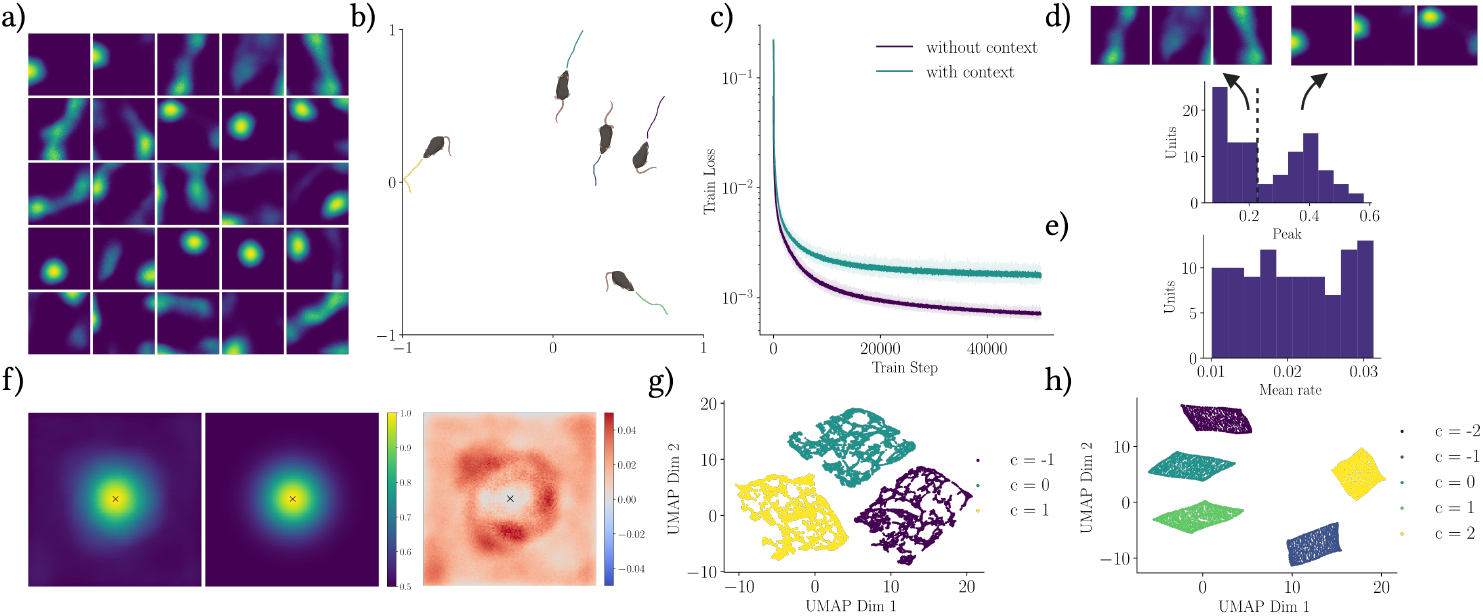
Recurrent network results with and without context. a) Ratemap examples of randomly selected units of a recurrent network without context. b) Example trajectories used for training. c) Training loss for recurrent networks with and without context (10 models each, error bands show min and max). d) Histogram of peak values of a recurrent network without context and example ratemaps of units of different parts of the distribution. e) Histogram of mean rates of a recurrent network without context. f) Similarity structure in the center location of the learned representations of a recurrent network without context (left) and the objective (center), as well as the difference between the two (right). g) First two UMAP dimensions of a selection of trajectories sampled from different contexts. h) First two UMAP dimensions of spatial representations in different contexts.

Besides being able to simply represent location and context conjunctively, we find that the recurrent network learns low-dimensional structures representing distinct contexts. By applying UMAP to recurrent representations, we observe that low-dimensional projections indeed capture both the geometry of the square enclosure, and the identity of the context, even when representations are aggregated from trajectories collected from multiple different contexts (Fig. 4g) and h)). It therefore appears that the network has learned veritable cognitive maps [O’Keefe and Nadel, 1978] of different contexts, which can be readily accessed from the representations themselves.

### 3.4 Reuse by rotation

Having demonstrated that both feedforward and recurrent networks learn remapping through distinct representations of different contexts, we now turn to an interesting feature of the similarity objective. Assuming that we have a trained (feedforward) network, with representations **p**(**x**_*t*_) (for all **x**_*t*_ in the domain), the loss function only depends on the norm of, and distance between population vectors. Thus, the objective is invariant to an orthogonal transformation *Q* of the representation. In other words, we can transform the entire set of population vectors (as long as the transformation preserves pairwise distances and pointwise norms), and still have a representation that minimizes (2).

In fact, we find that orthogonal transformations can be used to achieve some of the representational changes learned by the network through training. Specifically, we computed orthogonal transformations to match spatial representations across different contexts (see 2.5 for a description). Fig. 5a) shows that these transformations achieve significantly smaller errors than random orthogonal transformations. Furthermore, they preserve the similarity structure, and replicate the remapping behavior observed in the learned representations, as illustrated in Fig. 5b).

**Figure 5.**
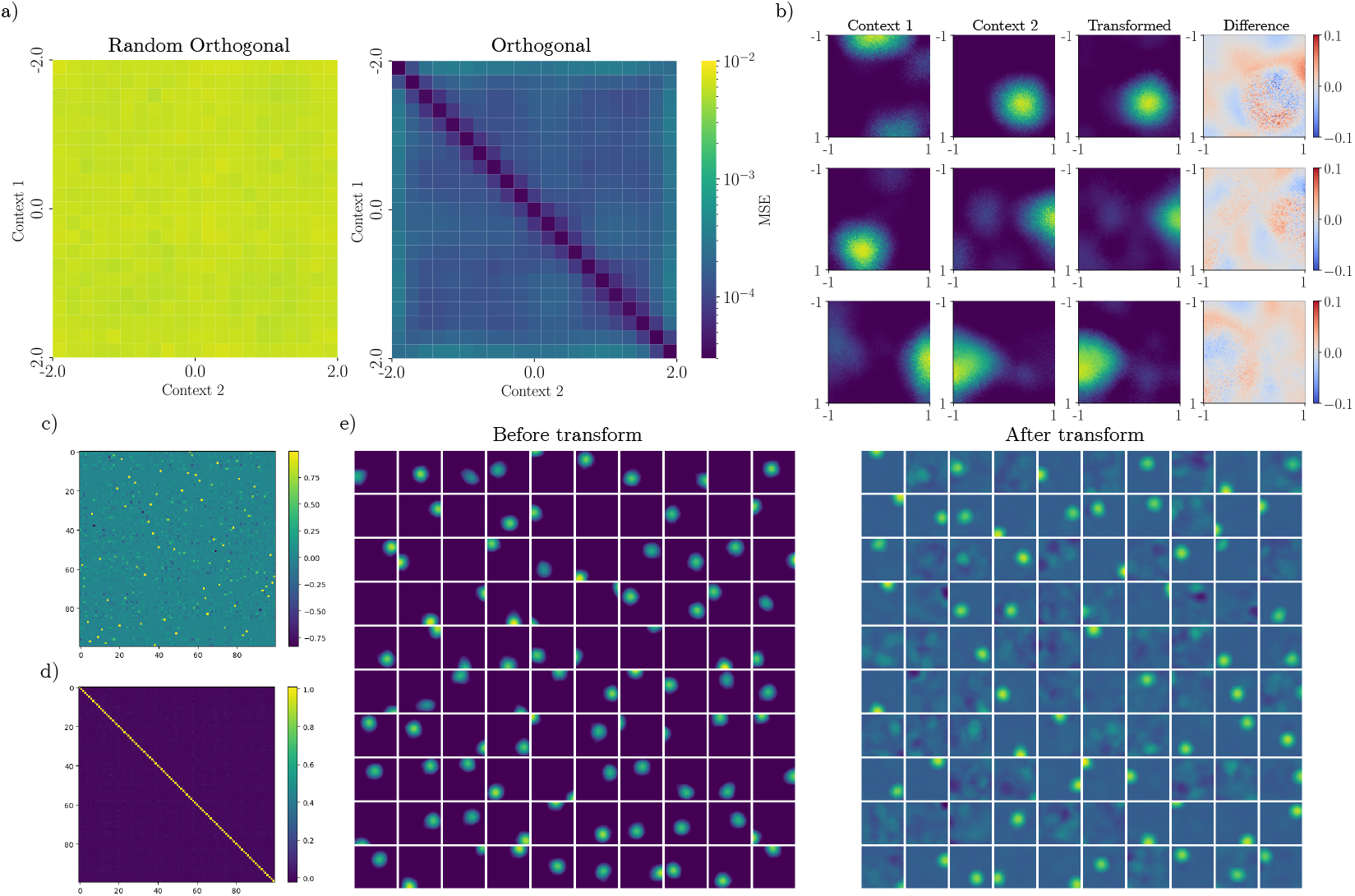
Remapping by orthogonal transformations. a) Mean squared error between learned representations of the recurrent network, and representations obtained by an orthogonal transformation, for different contexts. b) Example ratemaps of the recurrent network and corresponding transformed ratemaps, obtained using an orthogonal transformation. c) Elements of the learned, near-orthogonal matrix *Q*. d) The product *QQ*^*T*^. e) Feedforward network ratemaps before and after applying the near-orthogonal transformation *Q*.

Additionally, we trained a new orthogonal transformation (as described in 2.5) that attempted to preserve the non-negativity of the original representation (Fig. 5c) and d)). The effects of applying the transformation are shown in Fig. 5e), for the entire population. Notably, unit firing fields shift and morph, and some units turn on, or become effectively silenced, similar to remapping behavior.

As a result, restricted orthogonal transformations could prove a viable way of modeling Hippocampal remapping. These findings could also be extended to study what kind of upstream representations are needed to induce downstream representational rotations. Doing so could conceivably shed light on interactions between place cells and other cell types during remapping, in which other spatial cells such as grid cells are often implicated [Latuske et al., 2018].

### 3.5 Limitations

While our model provides a fresh perspective on the formation and remapping of Hippocampal cells, there are several factors that limit its scope. For one, we consider models where label locations and pre-processed context signals are available during training time, which could be biologically implausible. A related concern is the use of a scalar context signal. However, our model could accommodate more complex context signals, such as vectors with a range of different features, such as observational input. Extending the context signal in this way could prove to be an interesting avenue of research.

Another limitation is the use of rate coding directly. It is somewhat unclear how the proposed objective could be extended to spiking networks. A third limitation is the choice of the similarity lower bound, i.e. *β*. However, this could conceivably be determined by exploring representational similarity in experimental data in the future.

## 4 Conclusion

This work introduces a similarity-based objective to explain the functional characteristics of place cells. Using this minimal self-supervised objective, we are able to directly demonstrate how place cell-like representations can be learned, and how they can be understood as translating similarity in location, to similarity in representation. Furthermore, we observe emergent global remapping as a consequence of conjunctive encoding of space and contexts. Finally, we demonstrate that remapping may be enacted through orthogonal transformations, without the need to relearn the entire spatial environment. By demonstrating how place-like representations can emerge and adapt to different contexts, our findings contribute to a deeper understanding of the neural basis of navigation and memory.

